# On the application of BERT models for nanopore methylation detection

**DOI:** 10.1101/2021.02.08.430070

**Authors:** Yao-zhong Zhang, Sera Hatakeyama, Kiyoshi Yamaguchi, Yoichi Furukawa, Satoru Miyano, Rui Yamaguchi, Seiya Imoto

## Abstract

**Motivation:** DNA methylation is a common epigenetic modification, which is widely associated with various biological processes, such as gene expression, aging, and disease. Nanopore sequencing provides a promising methylation detection approach through monitoring abnormal signal shifts for detecting modified bases in target motif regions. Recently, model-based approaches, especially those with deep learning models, have achieved significant performance improvements on nanopore methylation detection. In this work, we explore using bidirectional encoder representations from transformers (BERT) for doing the task, which can provide non-recurrent neural structures for fast parallel computation.

**Results:** We find original BERT architecture does not work as well as the bidirectional recurrent neural network (biRNN) on the nanopore methylation prediction task. Through further analysis, we observe recurrent patterns of positional-signal-shift in the context window surrounding target 5-methylcytosine (5mC) and N6-methyladenine (6mA) motifs. We propose a refined BERT with relative position representation and center hidden units concatenation, which takes account of task-specific characters into modeling. We perform systematic evaluations in-sample and cross-sample. The experiment results show that the refined BERT model can achieve competitive or even better results than the state-of-the-art biRNN model, while the model inference speed is about 6x faster. Besides, on the cross-sample evaluation of datasets from the different research groups, BERT models demonstrate a good generalization performance.

**Availability:** The source code and data are available at https://github.com/yaozhong/methBERT

## 1 Introduction

Methylation of DNA/RNA/histone is commonly observed in developmental disorders, aging, and genomic disease, such as cancer. Fast and accurately detecting methylation status has a fundamental requirement to find distinctive biomarkers for aging/disease profiling. For a virome/metagenome study, quick and accurate epi-transcriptome detection also plays an important role in understanding unseen strains (Kim *et al*., 2020). One commonly used DNA methylation detection approach is Whole-Genome Bisulfite Sequencing (WGBS). To detect modified bases, WGBS first takes sodium bisulfite conversion before sequencing. As the pre-chemical bisulfite conversion is a relatively harsh process, it makes DNA sequences more fragmental and a large amount of DNA is usually required. Also, limited to the read length, it is difficult to align short reads in low-complex regions and analyze methylation patterns in a long-range. The data processing of WGBS is sophisticated and time-consuming. Various biases (e.g. GC and fragment length) including those introduced by bisulfite treatment are required to be dealt with in the data analysis. WGBS can only be used for DNA samples, which limits its application of detecting RNA methylation. Single-molecule sequencing (e.g., PacBio and Nanopore) provides a promising approach through detecting abnormal signals in target motif regions, as modified bases usually have different current signals. Compared with the sodium bisulfite approach, no extra chemical treatment is required, which helps to reduce potential biases.

Currently exist nanopore methylation detection methods can be categorized into two types. One is testing-based (e.g., Tombo (Stoiber *et al*., 2016)), the other is model-based (e.g., nanopolish (Simpson *et al*., 2017), deepMod(Liu *et al*., 2019) and deepSignal (Ni *et al*., 2019)). A testing-based approach performs statistical test on paired signals (candidate and reference) and does not require any training process. Also, it can be applied for any chemical modifications. A model-based approach trains a model on known chemical modifications and makes predictions whether a signal sequence contains methylation signals or not. Sequential models, such as hidden Markov model (HMM) and bidirectional recurrent neural network (biRNN), are commonly used in the model-based approach.

Although model-based approaches have already achieved competitive results, the sequential computational order makes them difficult to be optimized in parallel for fast inference. Meanwhile, finding discriminative signal patterns for identifying methylated signals is also important for developing novel detection algorithms. In this work, based on the bidirectional encoder representations from transformers (BERT), we explore the non-recurrent modeling approach for nanopore methylation detection. Though analyzing nucleotide sequences with both methylated and unmethylated signals, we profile positional signal-shift for different motifs and methyltransferases. We find ±3*bp* region surrounding the center methylation candidate shows significant signal-shifts. Different methylation types, such as 5-methylcytosine (5mC) and N6-methyladenine (6mA), also demonstrate different signal-shift patterns. We hence propose a refined BERT model to take account of signal-shift patterns in the modeling. We evaluate the proposed methods on the publicly available benchmark dataset. In both in-sample and cross-sample evaluation, the proposed refined BERT model achieves a competitive or even better result when compared with the state-of-the-art biRNN model, while its model inference speed is about 6x faster. In the cross-sample evaluation, BERT models also demonstrate their transfer learning ability across different datasets.

## 2 Methods

In this section, we introduce BERT (Devlin *et al*., 2018) and refined BERT applied for nanopore methylation detection. The BERT is built on the base of Transformer (Vaswani *et al*., 2017), which employs self-attention as the core module in its stacked network structure. It is proposed to replace recurrent and convolution operation with purely attention mechanisms. A typical transformer network consists of encoding and decoding module.

BERT only uses the encoding module of a typical transformer for pretraining on the unsupervised data. BERT has achieved break-through results on many natural language understanding tasks. In this work, we explore applying the BERT model for the nanopore methylation detection task to leverage the power of advanced deep learning models.

### 2.1 BERT and refined BERT model

Figure 1 shows the model structures of BERT models used for nanopore methylation detection. We explore two types of BERT models. One is the most commonly used BERT (Figure 1(a)), the other is the refined BERT (Figure 1(b)), which is optimized for nanopore methylation detection.

**Fig. 1:**
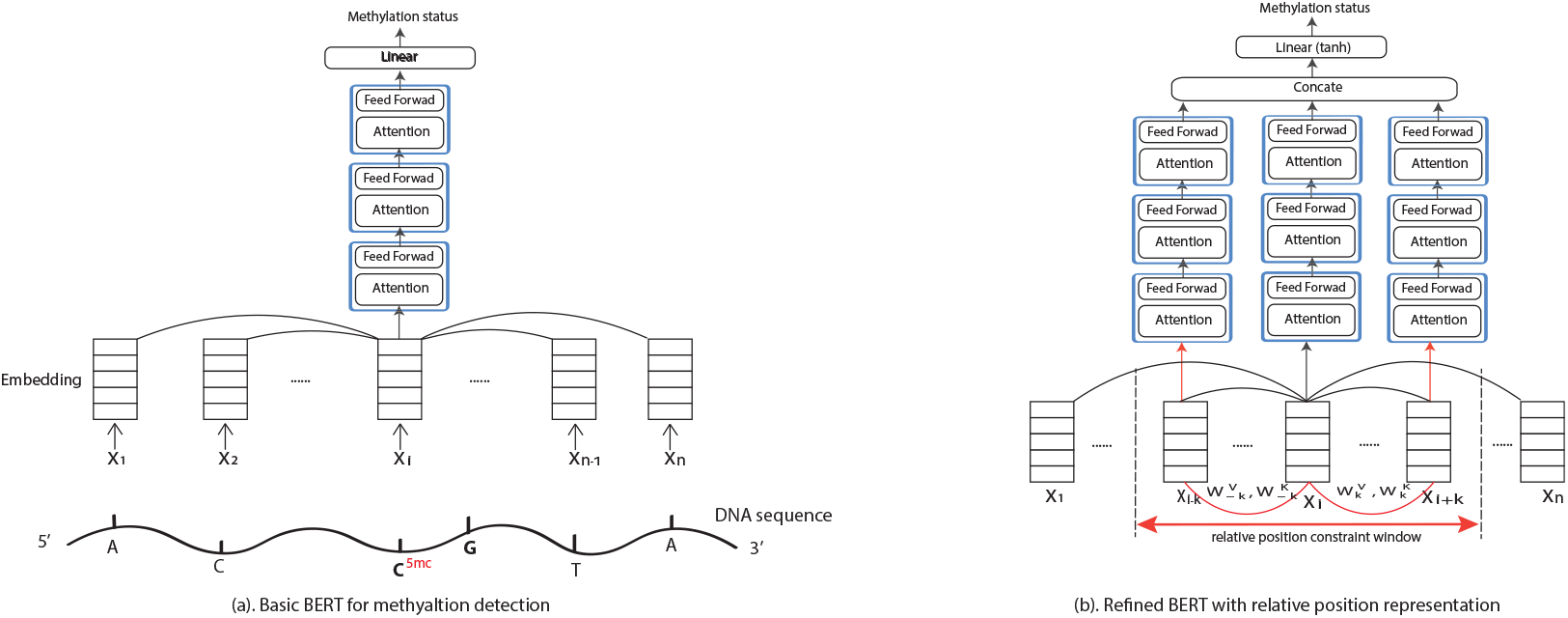
Basic BERT’s and refined BERT’s model structure used for methylation detection. Compared with the basic BERT, enhanced constraints and additional edges are highlighted in red color.

#### 2.1.1 Embedding module

Given extracted features for each position in a sequence, the embedding layer maps input vectors into hidden spaces. In the embedding layer, besides event embedding, positional embedding (PE) is also included. As a BERT is used to learn bidirectional contextual information, positional information is important in the modeling. The original PE (Vaswani *et al*., 2017) uses a sinusoid embedding, which is fixed and not learnable.

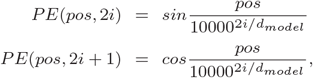

where *pos* is the position and *i* is the embedding dimension. For any fixed offset *k, PE_pos+k_* can be represented as a linear function of *PE_pos_*. According to the recent progress (Huang *et al*., 2020), learnable PE and relative position embedding can help to further improve BERT’s performances. Therefore, in the refined BERT model, we use learnable PE and relative position representation. The learnable PE takes positional embedding vectors as parameters, which are updated during the learning process.

#### 2.1.2 Self-attention module

Following the embedding layer, there are three stacked transformer blocks. Each transformer block consists of a multi-head self-attention layer and position-wise fully connected feed-forward network. The self-attention mechanism is a modeling approach of describing context information for different positions of inputs under a deep learning framework. The selfattention mechanism imitates the human sight mechanism and provides a model with the ability to zoom in or out in a particular position of an input sequence. It demonstrates the effectiveness in many different tasks including natural language understanding, image recognition, and several bioinformatics applications.

Attention function is described as mapping *Q* and a set of key-value (*K, V*) pairs to an output. Formally, for an input *x* = (*x_1_,…, x_n_*) of n elements where *x_i_* ∈ *R^d_x_^*, we calculate query *Q*, key *K* and value *V* vectors of dimension *d_k_* based on the embedding vector of *embed*(*x*). The attention module generates a new sequence *z* = (*z*_1_,…, *z_n_*) of the same length as *x*. *z_i_* is calculated as a weighted sum of linearly transformed input elements as follows:

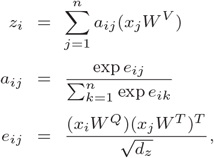

where *W^Q^, W^K^, W^T^* ∈ *R^d_x_×d_z_^* are parameter matrices.

The self-attention computes a pairwise correlation of *embed*(*x_i_*) and *embed*(*x_j_*), which can be calculated in a parallel way. While in a biRNN, recurrent hidden units are required to be calculated successively. This architecture difference makes BERT can be optimized for fast inference.

#### 2.1.3 Relative position representation in self-attention heads

For nanopore sequencing, signals are supposed to be more affected by the nucleotide passing through the pore. Its surrounding nucleotides may also have effects on the current signals. For those nucleotides that are too far away in a context window, it is intuitive to assume they have less effect on the detected current signals. In the refined BERT model, we add relative position representation in the attention module following the method proposed by Shaw et al. (2018). For any two input elements *x_i_* and *x_j_*, the relative position information is modeled with two distinct edge representations 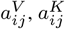. For linear sequences, those edges are used to capture the relative position differences between input elements. As the precise relative position is not useful beyond a certain distance, we clip the maximum distance (e.g. ±3bp) in calculating attention *a_ij_* ∈ *A*.

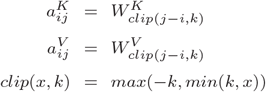

#### 2.1.4 Final full connection layer

After the stacked transformer blocks, hidden units of the center position feed to a full connection linear layer that makes the final prediction of whether a given input contains a methylated motif or not. In the refined BERT, besides the hidden units of the center position, hidden units in its surrounding window (e.g., ±3*bp*) are concatenated as the input of the final full connection layer.

### 2.2 Applying BERT models for nanopore methylation detection

The BERT models are then applied to replace different classification models (e.g. biRNN) in a typical model-based methylation detection framework. In this framework, raw signals of each read are first translated into nucleotide sequences (basecalling). Signals are then aligned to corresponding reference nucleotides through the re-squiggle process. After that, the target motif (e.g. CpG) and its context regions are localized through nucleotide matching and signals in a context window of a fixed length (e.g. 21bp) are transformed into event-based features as the input of methylation callers. Typical event-based features include signal mean, signal standard deviation, event length, and nucleotide information (Liu *et al*., 2019). Here, we utilize the framework of deepMOD and perform the same pre-process for the data. We use Tombo (Ver 1.5.1) to perform re-squiggling and utilize Minimap2 (Ver 2.17-r941) to align events to the reference genome. Here, we use E.coli K-12 MG1655 and H.Sapiens GRCh38 as the reference genomes.

## 3 Experiments

We compare BERT models with the state-of-the-art biRNN model, which is used as the basic network structure in DeepMOD (Liu *et al*., 2019) and DeepSignal (Ni *et al*., 2019). To compare with other non-deep-learning-based methods, we utilized the CpG benchmark pipeline (Yuen *et al*., 2020) as a pivot.

### 3.1 Data and model parameters

We train and test the models on the public accessible 5mC (Stoiber *et al*., 2016; Simpson *et al*., 2017) and 6mA (Stoiber *et al*., 2016) datasets. The datasets include samples of E.coli K-12 MG1655, K-12 ER2925, and H.sapiens NA12878. Negative control samples are amplified with PCR and no modified bases are included. Positive control samples are synthetically introduced by specific enzymes after PCR amplification, which includes SssI, Hhal, MpeI methylases for 5mC, and TaqI, EcoRI, and Dam for 6mA modification. We use the samples that are sequenced with Oxford Nanopore R9 flow cells. For each dataset, we randomly shuffle reads in positive and negative controls and construct the training, validate and test set according to a split proportion of 80/10/10 for in-sample evaluation. For the cross-sample evaluation, we train models on one dataset and test on the other dataset.

BiRNN uses the default model architecture and parameter setting of DeepMOD, which consists of three stacked bi-directional recurrent layers (hidden_size=100) and one full connection layer for the center position. The total number of biRNN parameters is 570,802 for an input length of 21bp. BERTs use three attention layers (hidden_size=100, attention_head=4) and one full connection layer. For the refined BERT, learnable positional encoding, attention with relative position representation and center-hidden-concatenation are used. For BERT and refined BERT, there are total of 364,902 and 368,202 parameters, which are around 35% less than that of biRNN. More detailed information on the model structures is described in the supplement material. We implement the three models using Pytorch. All the models are optimized using Adam optimizer (Kingma and Ba, 2014) with the learning rate of 1e − 4 and maximum iteration epoch of 50. Model parameters are selected based on the minimum validation loss.

### 3.2 Exploring differentiated signal positions in the context window surrounding target motifs

Ideally, we assume a modified nucleotide (e.g., the center position of XXXXXXXXXXC^5*mC*^ GXXXXXXXXX) has different current signals, when compared with the unmodified one. As the boundary of nucleotide/k-mer signals are not rigorous and surrounding nucleotides may also be affected, it is worthwhile investigating signal-shift patterns related to methylation in a large context. To identify signal-shift affected by methylation for a specific dataset, we use a simple quantification approach to calculate significant signal changes of each position in the context window. Given a dataset of a specific motif and methyltransferase, we first cluster instances with the same nucleotide sequence to avoid the effect of nucleotide sequences. We reserve sequence clusters that contain both methylation and unmethylation instances (≥ 1). For each sequence cluster, we normalize event signal values of methylation samples with their according unmodified averaged event signal values for each position. The *i*-th positional signal-shift is then calculated as 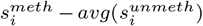. For those normalized methylation samples, we calculate basic statistics of signal-shift for each position and draw boxplots for 5mC and 6mA training sets.

Shown in Figure 2, for all datasets, we can observed positions of significantly signal-shift are located in a range of ±3bp to the center position (the 11th) in which the target nucleotide is located. For the rest off-center positions, the averaged signal-shift values are close to 0. This indicates a modified nucleotide not only affect its corresponding current signals but also the signals of its surrounding nucleotides.

**Fig. 2:**
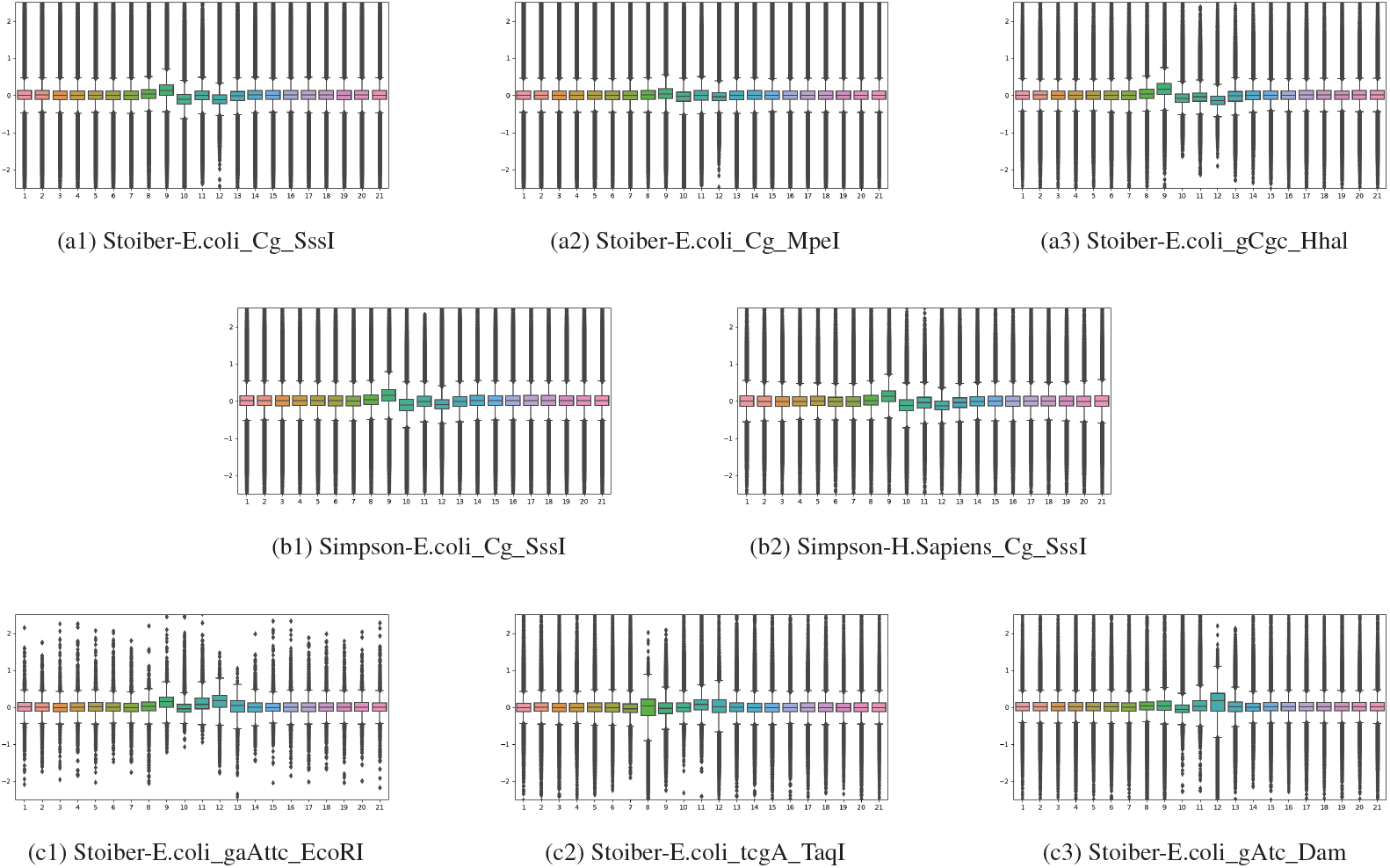
Boxplot of positional signal-shift for 5mC and 6mA datasets of the specific motif and methyltransferase. (a1),(a2) and (a3) are on Stoiber’s E.coli 5mC dataset. (b1) and (b2) are on Simpson’s 5mC dataset. (c1), (c2) and (c3) are on Stoiber’s E.coli 6mA dataset. Each dataset is represented in a format of dataSource_motif_methltansferase.

Besides, 5mC and 6mA datasets show different positional-signal-shift patterns. Specific positions, such as −2bp position (9th) in the 5mC dataset and +1bp position (12th) in the 6mA dataset, have larger averaged signal-shift values. Such pattern can be generalized across the different dataset with the same motif and methyltransferase. For example, Figure 2 (a1), (b1) and (b2) show a similar positional signal-shift pattern. For different methyltransferases, such as Hhal (Figure 2(a3)) also shows a similar pattern as in SssI, while MpeI does not have a similar pattern obviously (Figure 2(a2)).

Those positional signal patterns can be directly modeled by a biRNN, while for the basic BERT, they are not specifically considered in its model structure. In a biRNN, such as the implementation of deepMOD, the last full connection layer uses hidden units of the center time step as the input. Meanwhile, the bi-directional structure and the information decay from both ends to the center position render the model focusing more on center positions. For the basic BERT, as any arbitrary timestep pair is processed with the same attention module, the importance of center positions are not specifically considered in the model. Therefore, we propose a refined BERT model to solve this problem. We incorporate relative-position attention and center-hidden-units concatenation to enable a BERT model to pay more attention to center positions.

### 3.3 In-sample evaluation

To evaluate model performance, we first perform the in-sample evaluation on 5mC and 6mA datasets. The predictions of different models are evaluated on the read and genomic level. For the genomic level evaluation, we group all reads aligned to the same genomic coordinate, and uses a threshold of prediction methylation percentage ≥ 0.1 (same as deepMOD) as a genomic position prediction.

In general, on the five 5mC datasets, the AUC performance of the three models are relatively close on both read level and genomic level. The basic BERT model does not work as well as the biRNN model that AUC scores are lower. The refined BERT model achieves equivalent or better AUC scores on the genomic-level. Note that on the dataset *Stoiber_E.coli_CG_Mpel* and *Simpson_E.coli_CG_SssI*, although the read-level AUC of the refined BERT are 0.0014 and 0.005 lower than that of biRNN, the genomic-level performance of the refined BERT is equal or significantly better than biRNN. This can be explained by the more accurate prediction in several low read-coverage regions. On the 6mA dataset, the refined BERT model achieves the best AUC performance on both read-level and genomic-level. The performance of the basic BERT model is variant and unstable. On *Stobier_E.coli_gaAttc_EcoRI* and *Stoiber_E.coli_gAtc_Dam*, the basic BERT performs slightly better than biRNN on the read-level AUC, but has a large performance gap on *Stoiber _E.coli _gaAttc_EcoRI*.

In summary, in the in-sample evaluation, the refined BERT model can achieve competitive or better results when compared with the biRNN model on benchmark 5mC and 6mA datasets.

### 3.4 Cross-sample evaluation

We then conduct the cross-sample evaluation. To compare with other non-deep-learning based methods, we utilize the benchmark pipeline (Yuen *et al*., 2020) as a pivot. We test models on the same benchmark dataset1, which is generated based on Simpson’s E.coli dataset with different methylation levels. In the dataset, 100 arbitrary sites are selected, which contain singleton CpG in a window of 10nt from both methylated and unmethylated instances in the Simpson’s E.coli dataset. Yuen et al. created 11 specific mixtures of methylated and unmethylated reads, containing 0%, 10%,…, 100% of methylated reads. Each mixture contains approximately 2400 reads. More detailed information can be found in (Yuen *et al*., 2020).

Different from the deepMOD model used in the original benchmark pipeline, which is pre-trained on a mixture dataset of all 5mC positive (Cg_SssI, Cg_MpeI, and gCgc_Hhal) and negative controls (UMR, con1, and con2). Here, we test two different models trained on a single dataset with the same methyltransferase to reduce potential overlapping between the training and testing set. All three models are trained on *Stoiber_Ecoli_CG_SssI* and *Simpson_Hsapiens_CG_SssI*, separately. *Simpson_H sapiens_CG_SssI* is sequenced by the same group on different species, while *Stoiber*_*Ecoli*_*CG*_*SssI* is sequenced by a different group on the same species. We use METEORE pipeline (Yuen *et al*., 2020) to generate violin plots for model predictions on each mixture. The Pearson’s correlation *r*, coefficient of determination *r*^2^ and root mean square error (RMSE) are used as the evaluation metrics for each model.

With the training data of *Simpson_Hsapiens_CG_SssI*, all three models achieve performances ranked next to the best reported results of Megalodon(r=0.9860, *r*^2^ = 0.9723, RMSE=0.0758) on the dataset (Yuen *et al*., 2020). BiRNN achieves the best Pearson correlation r=0.9828 and *r*^2^=0.9658, while refine BERT achieves minimal RMSE of 0.0732 among the evaluated three models.

When using *Stoiber*_*Ecoli*_*CG*_*SssI* for training models, the performances of all three models decrease. This indicates the challenge of using datasets sequenced by different research groups. Here, both BERT models show better performances than biRNN, as in Figure 3b. The refined BERT achieves the best r=0.9446, *r*^2^ =0.8924 and RMSE of 0.1449 among the three models, which demonstrate the generalization ability on datasets sequenced by different research groups. Based on the reported benchmark results, the Pearson correlation ranks between reported deepMOD and deepSignal (Megalodon > DeepMOD_*mixModel*_ (0.9467) > refined BERT > DeepSignal_*human_hx1*_ (0.9420) >Guppy>Nanopolish>Tombo).

**Fig. 3:**
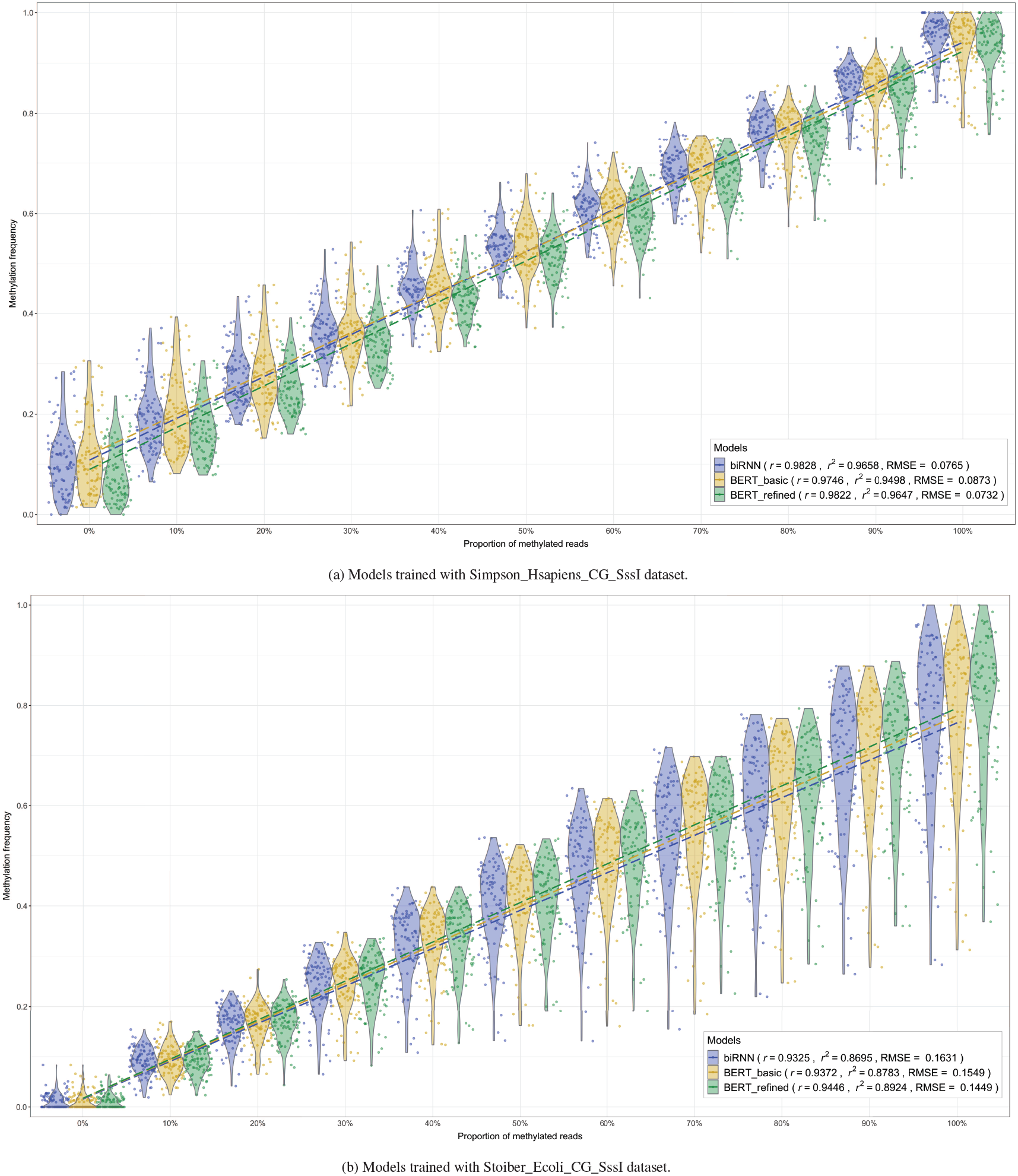
Violin plots of prediction results of models trained on different datasets.

### 3.5 Model inference speed

The main motivation of applying BERT models is to use a non-recurrent modeling approach for the nanopore methylation detection task to improve the model inference speed. We performed a speed test on a server with 24 CPU cores (Intel(R) Xeon(R) Gold 6126 CPU @ 2.60GHz) and one V100 NIVIDA GPU card. In the running, CPUs are responsible for data loading and feature extraction, while GPU works for model inference. We tested the model inference time and total running time of the three models on the benchmark dataset1. For each mixture split, we repeated 5 times running and took the averaged value. As shown in Table 3, the model inference speed of BERT models is around 6x~7x faster than biRNN model (BERT_refined:5.96x, BERT_basic:7.16x). The inference time of refined BERT is only slightly slower than the basic BERT model. The gap of the total time is not that large (BERT_refined:1.14x, BERT_basic:1.16x), as the data I/O and feature extraction take major time. In the current implementation of BERT, we use reads as the basic data unit and integrate the data pre-processing part during a read-batch loading process. The data I/O and feature extraction part can be further accelerated.

**Table 1.**
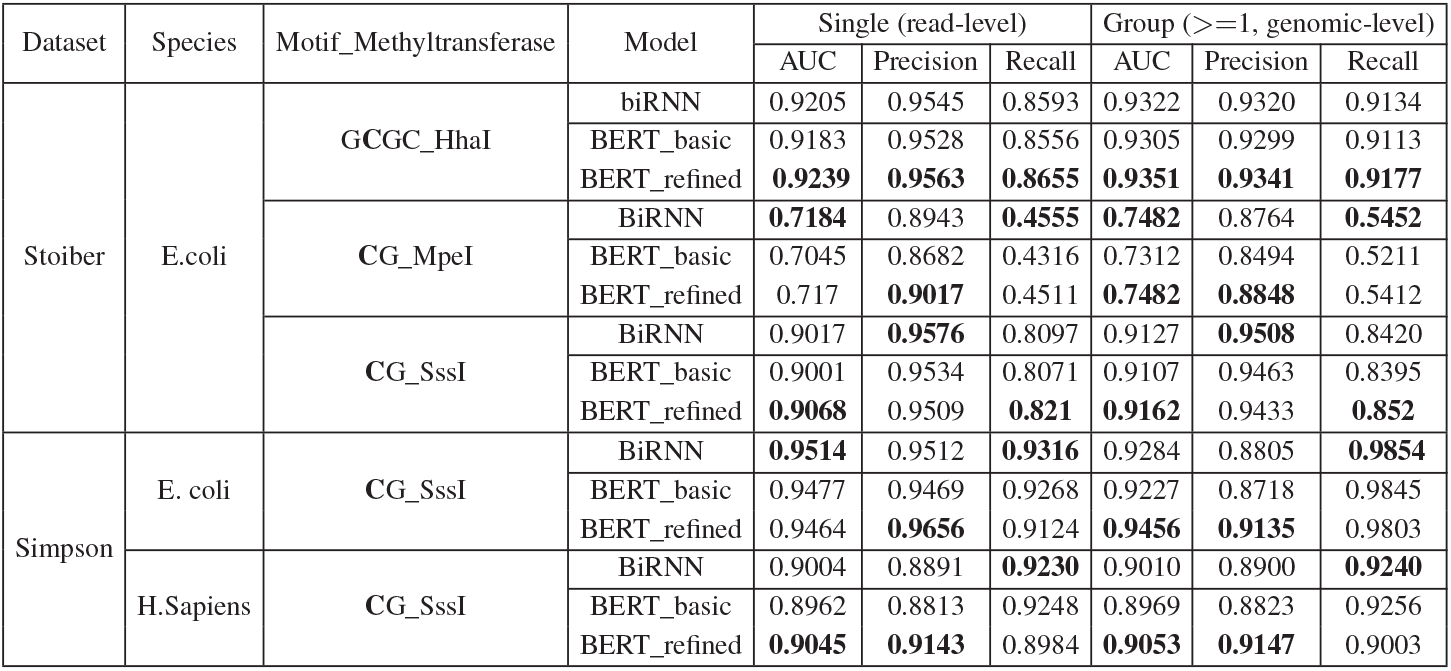
In-sample evaluation of different deep learning models on 5mC datasets. The best score of each dataset is highlighted in bold.

**Table 2.** In-sample evaluation of different deep learning models on 6mA datasets.The best score of each dataset is highlighted in bold.

| Dataset | Species | Motif_Methyltransferase | Model | Single (read-level) | | | Group ($\geq 1$, genomic level) | | |
| --- | --- | --- | --- | --- | --- | --- | --- | --- | --- |
|  |  |  |  | AUC | Precision | Recall | AUC | Precision | Recall |
| Stoiber | E.coli | gaAttc_EcoRI | BiRNN | 0.8524 | 0.8088 | 0.7497 | 0.8429 | 0.7797 | 0.8035 |
|  |  |  | BERT_basic | 0.8607 | 0.8151 | **0.7653** | 0.8591 | 0.7969 | **0.8277** |
|  |  |  | BERT_refined | **0.8611** | **0.8826** | 0.7473 | **0.8655** | **0.8596** | 0.7987 |
|  |  | tcgA_TaqI | BiRNN | 0.7722 | 0.7922 | 0.5750 | 0.7750 | 0.7789 | 0.6290 |
|  |  |  | BERT_basic | 0.7573 | **0.8168** | 0.5392 | 0.7653 | **0.8063** | 0.5937 |
|  |  |  | BERT_refined | **0.7857** | 0.7788 | **0.6064** | **0.7843** | 0.7643 | **0.6586** |
|  |  | gAtc_Dam | BiRNN | 0.6123 | **0.7656** | 0.247 | 0.6337 | **0.7631** | 0.3241 |
|  |  |  | BERT_basic | 0.6128 | 0.7329 | 0.2529 | 0.631 | 0.7311 | 0.3305 |
|  |  |  | BERT_refined | **0.6188** | 0.7513 | **0.2634** | **0.6385** | 0.7471 | **0.3421** |

**Table 3.**
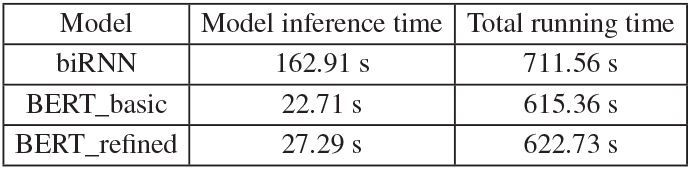
Model inference and total running time on the benchmark dataset1 for all 26402 reads.

## 4 Discussion

A BERT commonly works in a pre-training and fine-tuning approach. In the pre-training phase, a BERT learns bi-directional representations from unlabeled data. After that, learned feature representations are used on task-specific data for further fine-tuning. It has lead to several state-of-the-art results on many downstream tasks in language understanding. According to the data scale, the number of BERT parameters is usually large, and training such a model requires a huge amount of computational resources. For example, the BERT used for natural language modeling has a parameter scale ranging from 110M to 340M (Devlin *et al*., 2018). In this work, we did not follow this schema. Instead, we utilized the model architecture of BERT to provide a lightweight and non-recurrent solution to replace the recurrent biRNN model. In our experiment, the BERT uses three attention layers with 4 attention heads and 100 hidden units for each layer. The total number of model parameters is around 0.37M, which is even less than that of biRNN (0.57M). In the future, when more nanopore methylation data becomes available, a larger BERT model and pre-training and fine-tuning scheme can be further explored.

## 5 Conclusion

In this work, we explored applying BERT models for nanopore methylation detection, which aims to use a non-recurrent modeling approach for fast inference. We quantified positional signal-shift related to methylation for different datasets of specific motif/methylase and found patterns across datasets. In the process of evaluation, we found the original BERT architecture does not work as well as biRNN. We proposed a refined BERT considering task-specific characters into the modeling. Compared with the original BERT, the refined BERT uses learnable positional encoding and self-attention with relative position representation, and focuses more on the center positions in a ±3bp range. The experiment results show that the refined BERT can achieve competitive and even better results than the state-of-the-art biRNN model on a set of 5mC and 6mA benchmark datasets, while the model inference speed is about 6x faster. On the cross-sample evaluation, for the case that train and test data from different research groups, BERTs (include the original BERT) show a better performance than biRNN.

## Supporting information

More detailed information on the model structures is described in the supplement material

## Acknowledgements

We would like to thank Marcus Stoiber and Jared Simpson for making nanopore methylation data publicly available, Zaka Wing-Sze Yuen for providing the benchmark dataset and pipeline, authors of deepMOD and deepSignal for providing their source codes.

