## Supplementary material for "On the application of BERT models for nanopore methylation detection": More detailed information on the model structures is described in the supplement material

### Supplement materials

Network structures of related deep learning models

#### 1.BERT_basic

```
==========================================================================================
Layer (type:depth-idx)                   Output Shape              Param #
==========================================================================================
├──BERTEmbedding: 1-1                    [-1, 21, 100]             --
|    └──Linear: 2-1                      [-1, 21, 100]             800
|    └──PositionalEmbedding: 2-2         [-1, 21, 100]             --
|    └──Dropout: 2-3                     [-1, 21, 100]             --
├──ModuleList: 1                         []                        --
|    └──TransformerBlock: 2              []                        --
|    |    └──SublayerConnection: 3-1     [-1, 21, 100]             200
|    |    └──SublayerConnection: 3-2     [-1, 21, 100]             200
|    |    └──PositionwiseFeedForward: 3-3  [-1, 21, 100]           80,500
|    |    └──Dropout: 3-4                [-1, 21, 100]             --
|    └──TransformerBlock: 2              []                        --
|    |    └──SublayerConnection: 3-5     [-1, 21, 100]             200
|    |    └──SublayerConnection: 3-6     [-1, 21, 100]             200
|    |    └──PositionwiseFeedForward: 3-7  [-1, 21, 100]           80,500
|    |    └──Dropout: 3-8                [-1, 21, 100]             --
|    └──TransformerBlock: 2              []                        --
|    |    └──SublayerConnection: 3-9     [-1, 21, 100]             200
|    |    └──SublayerConnection: 3-10    [-1, 21, 100]             200
|    |    └──PositionwiseFeedForward: 3-11  [-1, 21, 100]          80,500
|    |    └──Dropout: 3-12               [-1, 21, 100]             --
├──Linear: 1-2                           [-1, 2]                   202
==========================================================================================
Total params: 243,702
Trainable params: 243,702
Non-trainable params: 0
Total mult-adds (M): 0.51
==========================================================================================
Input size (MB): 0.00
Forward/backward pass size (MB): 0.40
Params size (MB): 0.93
Estimated Total Size (MB): 1.33
==========================================================================================
 |- Total Model Parameters: 364902
```

#### 2.BERT_refined

```
==========================================================================================
Layout (type:depth-idx)                    Output Shape              Param #
==========================================================================================
├──BERTEmbedding_plus: 1-1                 [-1, 21, 100]             --
|    └──Linear: 2-1                        [-1, 21, 100]             800
|    └──PositionalEmbedding_plus: 2-2      [-1, 21, 100]             2,100
|    └──Dropout: 2-3                       [-1, 21, 100]             --
├──ModuleList: 1                           []                        --
|    └──TransformerBlock_relative: 2       []                        --
|    |    └──SublayerConnection: 3-1       [-1, 21, 100]             200
|    |    └──SublayerConnection: 3-2       [-1, 21, 100]             200
|    |    └──PositionwiseFeedForward: 3-3  [-1, 21, 100]             80,500
|    |    └──Dropout: 3-4                  [-1, 21, 100]             --
|    └──TransformerBlock_relative: 2       []                        --
|    |    └──SublayerConnection: 3-5       [-1, 21, 100]             200
|    |    └──SublayerConnection: 3-6       [-1, 21, 100]             200
|    |    └──PositionwiseFeedForward: 3-7  [-1, 21, 100]             80,500
|    |    └──Dropout: 3-8                  [-1, 21, 100]             --
|    └──TransformerBlock_relative: 2       []                        --
|    |    └──SublayerConnection: 3-9       [-1, 21, 100]             200
|    |    └──SublayerConnection: 3-10      [-1, 21, 100]             200
|    |    └──PositionwiseFeedForward: 3-11 [-1, 21, 100]             80,500
|    |    └──Dropout: 3-12                 [-1, 21, 100]             --
├──Linear: 1-2                             [-1, 2]                   1,402
├──Tanh: 1-3                               [-1, 2]                   --
==========================================================================================
Total params: 247,002
Trainable params: 247,002
Non-trainable params: 0
Total mult-adds (M): 0.51
==========================================================================================
Input size (MB): 0.00
Forward/backward pass size (MB): 0.42
Params size (MB): 0.94
Estimated Total Size (MB): 1.36
==========================================================================================
 |- Total Model Parameters: 368202
```

#### 3. biRNN model

```
biRNN(
  (lstm): LSTM(7, 100, num_layers=3, batch_first=True, bidirectional=True)
  (fc): Linear(in_features=200, out_features=2, bias=True)
)
 |- Total Model Parameters: 570802
```
